# A pan-cnidarian microRNA is an ancient biogenesis regulator of stinging cells

**DOI:** 10.1101/2022.12.15.520629

**Authors:** Arie Fridrich, Miguel Salinas-Saaverda, Itamar Kozlolvski, Joachim M Surm, Eleni Chrysostomou, Abhinandan M Tripathi, Uri Frank, Yehu Moran

## Abstract

An ancient evolutionary innovation of a novel cell-type, the stinging cell (cnidocyte), appeared >600 million years ago in the phylum Cnidaria (sea anemones, corals, hydroids, and jellyfish). A complex bursting nano-injector of venom, the cnidocyst, is embedded in cnidocytes and enables cnidarians paralyzing prey and predators, contributing to this phylum’s evolutionary success. In this work, we show that post-transcriptional regulation by a pan-cnidarian microRNA, miR-2022, is essential for biogenesis of these cells. By manipulation of miR-2022 levels in a transgenic reporter line of cnidocytes in the sea anemone *Nematostella vectensis*, followed by transcriptomics, single-cell data analysis, prey paralysis assays, and cell sorting of transgenic cnidocytes, we reveal that miR-2022 enables cnidocyte biogenesis, while exhibiting a conserved expression domain with its targets in cnidocytes of other cnidarian species. Thus, here we reveal one of nature’s most ancient microRNA-regulated processes by studying the functional basis for its conservation.

## Introduction

microRNAs (miRNAs) are small non-coding RNAs of 20-24 nucleotides that regulate messenger RNA (mRNA) levels in plants and animals^1–3^. In bilaterians (animals with bilateral symmetry such as arthropods, nematodes and vertebrates) these molecules are loaded into Argonaute (AGOs) proteins and bind mRNAs through a partial base-pair complementarity^4^ (~7 nucleotides at position 2-8 of the miRNA, known as “seed”). Consequently, the RNA Induced Silencing Complex (RISC) and the AGOs at their core are guided by the miRNA to reduce target mRNA levels as well as promoting translational inhibition^2^. This enables the regulation of cell differentiation and developmental processes. Among animals, the importance of individual miRNAs has been demonstrated in common lab model organisms belonging to the Bilateria group^5–8^.

Cnidarias (sea anemones, corals, hydroids and jellyfish) comprise a sister group that evolved in parallel to bilaterians for over 600 million years with a different set of their own microRNAs. Cnidarian miRNAs exhibit an ancestral mode of action as they bind their mRNA target with high complementarity (similarly to plants) to mediate target cleavage by the catalytic domain of AGOs^9^. Additionally, homologs of an ancestral miRNA biogenesis factor which was considered plants specific: Hyponastic Leaves 1 (Hyl1), were found in cnidarians but not in bilaterians^10,11^. miRNA turnover is higher in cnidarians relatively to bilaterians9,12. Yet, miR-2022 is one of the most conserved metazoan miRNAs and is found in all genomes of cnidarians sequenced to date^9,12–16^. Thus, it is one of only 2 pan-cnidarian miRNAs. However, miR-2022 function remained unknown since its discovery in 2008^12^. In the sea anemone *Nematostella vectensis*, miR-2022 is expressed in cnidocytes^9^.

Cnidocytes are a cnidarian defensive/offensive cell-type innovation that originated from the neurosecretory lineage^17–19^ >600 million years ago (mya) after the divergence from Bilateria. This cell lineage is essential for members of this ancient phylum to capture prey and repel predators through biogenesis of the cnidocyst: a stinging organelle that explosively discharges and either entangles or injects the prey or predator with venom^20,21^. The cnidocyst is one of the most complex biological structures in nature and its discharge is one of the fastest known biomechanical processes^22–24^.

While extensive efforts were devoted to identification of structural and venomous components of these organelles^20,25–32^, little is known about the regulatory mechanisms that enable their biogenesis beyond transcription factors. Nematogalectin Related 2 (NR2) is a structural component of the nematocyst tubule that is targeted by and co-expressed with miR-2022 in *Nematostella^9^*.

Notably, the functional role of individual miRNAs in animals was never explored in non-bilaterians. Thus, in this work, we tested the role of miR-2022 and the functional basis for its conservation across 600 million years of cnidarian evolution.

## Results and Discussion

### miR-2022 contributes to biogenesis of cnidocytes and fitness of Nematostella

miRNA precursors are stem-loop structures that are recognized by miRNA biogenesis factors such as Dicer and Drosha^33^. To test if Nv-miR-2022 takes part in the biogenesis of cnidocytes we designed a morpholino antisense oligo (MO) to interfere with Nv-miR-2022 biogenesis by binding the pri-miR and hence masking Dicer and Drosha processing sites^34^ (**Figure S1A**). Knockdown specificity was validated using stem-loop real time quantitative PCR (RT-qPCR)^35^ four days post injection of the MO to *Nematostella* zygotes **(Figure 1A)**. Wild type females were crossed with a transgenic male that expressed the fluorescent protein memOrange2 under the regulation of the promoter of the cnidocyte specific gene *NvNcol-3* (Ncol3::memOrange2^28^). As a result, ~50% of the progeny exhibited fluorescently labeled cnidocytes **(Figure 1B**, left panel). After injecting the zygotes of WT/Ncol3::memOrange2 cross, with Nv-miR-2022-MO we visually detected an overall reduction of fluorescent cnidocytes **(Figure 1B).** While the overall percentage of cnidocytes in *Nematostella* is low during the first few days of development (~2% of all cells^36^), we could easily detect cnidocyst capsules among dissociated cells of control injected animals in contrast to animals injected with miR-2022-MO **(Figure S1B).** Next, dissociated cells of miR-2022 morphants were subjected to flow cytometry analysis which enabled us to quantify a significant reduction in the number of fluorescently labeled cnidocytes **(Figure 1C–D, Figure S1C-D).** Notably, the fraction of highly fluorescent cnidocytes was more reduced than the fraction composed of weaker expression **(Figure S1C).**

**Figure 1:**
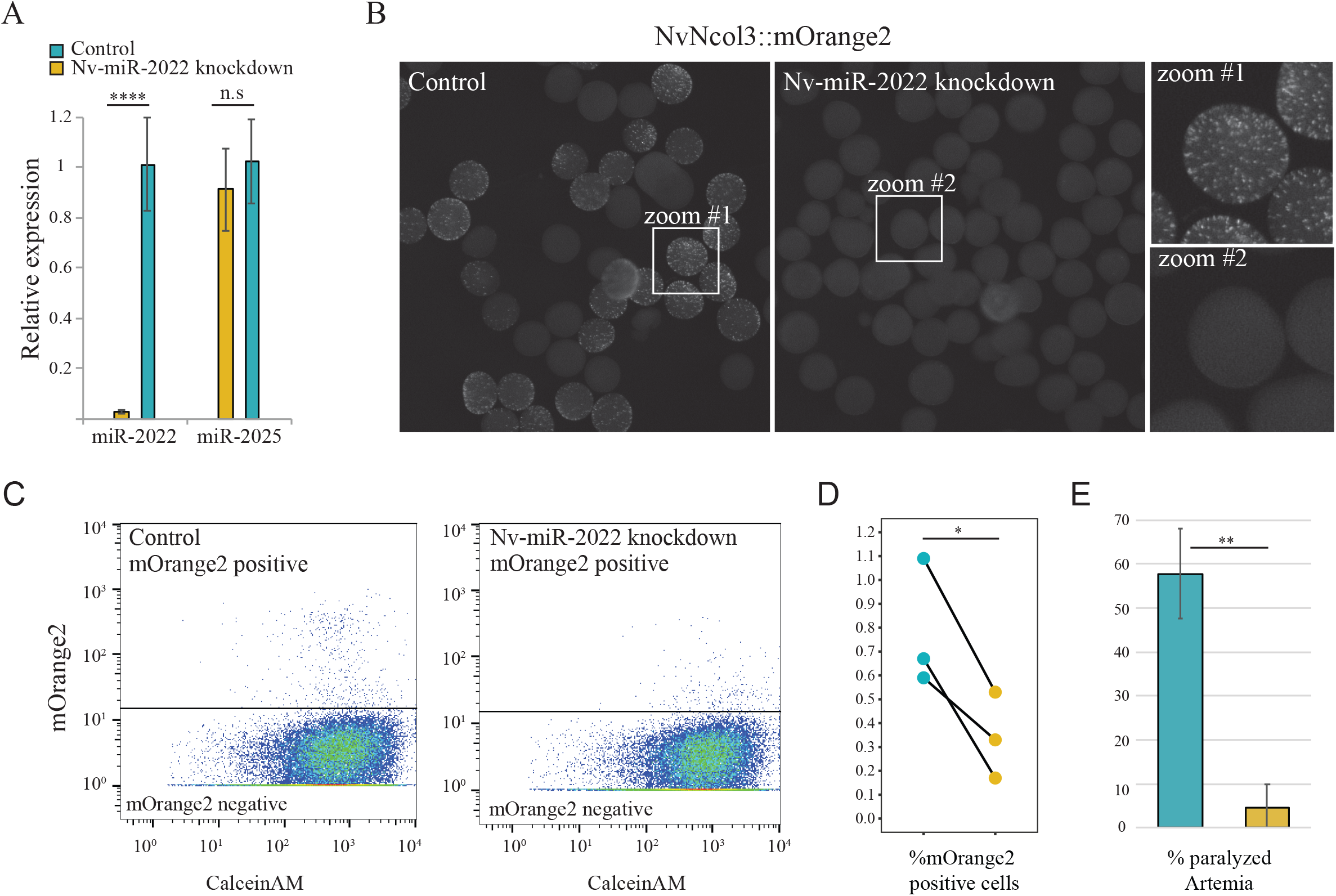
The impact of miR-2022 knockdown on cnidogenesis and prey paralysis potency in *Nematostella*. **(A)** Stem-loop qPCR validation of Nv-miR-2022 levels in *Nematostella* planulae 4 days post injection of control or miR-2022 MO into zygotes. Three independents biological replicates. (B) Intensity of mOrange2 in progeny of WT crossed with mOrange2::Ncol3 transgenes 4 days post injecting the zygotes with control or miR-2022 MO. Three independent biological replicates were subjected to flow cytometry analysis and the percentage of mOrange2 positive cells was quantified **(C-D). (E)** Percentage of paralyzed Artemia 5.5 hours post incubation with control or miR-2022 MO 4 days old *Nematostella* planulae (Three independent biological replicates).

We have previously shown that global inhibition of the miRNA pathway by knocking down key biogenesis factors in *Nematostella* results in severe developmental defects which prevents the metamorphosis from the swimming planulae stage to sessile polyps^11,37,38^. However, the knockdown of miR-2022 did not impact this developmental transition and 9 days post injection we observed metamorphosed polyps **(Figure S1E).** Thus, we conclude that miR-2022 plays a vital and specific role in the regulation of cnidocyte biogenesis.

*Nematostella* encodes two paralogs of AGO proteins: NvAGO1 and NvAGO2. Interestingly, while NvAGO1 can be found in diverse cell types, it exhibits a strong enrichment in the cluster that corresponds to cnidocytes in *Nematostella* single-cell RNA sequencing data^19^ **(Figure S1F).** Using NveAGO1 and NveAGO2 immunoprecipitations, we previously showed that miR-2022 exhibits the highest preference to load into NvAGOl^37^, demonstrating the overall regulatory importance of the miRNA pathway in cnidocytes.

Next, we hypothesized that the defensive capabilities of morphants of Nv-miR-2022 would be reduced due to the reduction in mature cnidocytes. Previously we showed the defensive capability of young *Nematostella* larvae through stinging potential predators^27^. We utilized a similar assay and incubated 4-days-old *Nematostella* planulae, which were injected with either control or Nv-miR-2022 MO, with nauplii of the brine shrimp *Artemia salina.* Five and a half hours post-incubation, *Artemia* incubated with control injected *Nematostella* showed a significantly higher number of paralyzed/dead individuals compared to *Artemia* incubated with Nv-miR-2022 MO injected *Nematostella* (*t*-test, p=0.009489, **Figure 1E**, video **Data S1).**

The loss of cnidocytes in *Nematostella* miR-2022 morphants as well as their reduced potency in paralyzing *Artemia*, yet without impacting the developmental metamorphosis of the animal, suggest that miR-2022 carries a specific role to regulate a restricted cell-type population that is vital for the survival of cnidarians.

To further explore the specificity of miR-2022 function in cnidocytes we measured gene expression levels in *Nematostella* miR-2022 morphants.

### Transcriptome of *Nematostella* miR-2022 morphants is enriched with mis-regulated cnidocyte-specific genes

To assay the effect of Nv-miR-2022 knockdown on gene expression we extracted RNA from 4 days old Nv-miR-2022 morphants and animals injected with control MO and employed high-throughput RNA sequencing. We carried out differential expression analysis using DESeq2^39^ which enabled us to identify 944 differentially expressed genes (DEGs). The majority of DEGs in miR-2022 morphants appeared to be down regulated (805 of 944) (**Figure 2A**). Recently, our NvNcol3::memOrange2 transgenic line was utilized to isolate the cnidocytes of *Nematostella* primary polyps and profile their transcriptome^40^. We re-analyzed these transcriptomes and generated a list of 2,615 genes enriched in Ncol3::mOrange2 positive cells (**Figure S2, Data S2**). We found that 268 out of the 944 DEGs in miR-2022 morphants overlapped with the list genes enriched in Ncol3::mOrange2 positive cells (**Figure 2B**). To test if such an overlap represented a significant enrichment, we simulated 10,000 lists of randomly chosen 944 genes from *Nematostella* transcriptomes, and tested the probability of getting as many as 268 genes overlapping with the list of cnidocyte-specific genes by chance (see methods). We found an average peak for 109.54 overlapping genes per list, with not even a single list containing as many as 268 cnidocyte-enriched genes (permutation test, p=0, **Figure 2C).** Next, we analyzed a recently published *Nematostella* single cell dataset^19^ and examined which cell clusters contain the differentially expressed genes in Nv-miR-2022 morphants (**Figure 2D**). To this end, we generated a list of marker genes for each cell cluster resulting in 2168 marker genes that are distributed between the 12 cell clusters. We collected all Nv-miR-2022 morphant differentially expressed genes that overlapped with the single-cell cluster markers and found that 65.4% of these genes (110/168) were in clusters 1 and 2 that correspond to cnidocytes, while the rest were spread between clusters 3-12. (**Figure 2D**). The majority of these differential expressed genes were in cluster 2 which corresponds to mature cnidocytes. To test the significance of this enrichment we simulated 10,000 lists of random gene collections and never detected as many as 110 genes to overlap with cnidocyte clusters (permutation test, p=0, average peak = 18.32 genes per list, **Figure 2E**). The downregulation of many cnidocyte-specific genes in our miR-2022 morphants (**Figure 2**) is in line with our observation of reduction of Ncol3::memOrange2 positive cells in these animals (**Figure 1**), further supporting the involvement of miR-2022 in regulation of cnidocyte biogenesis.

**Figure 2:**
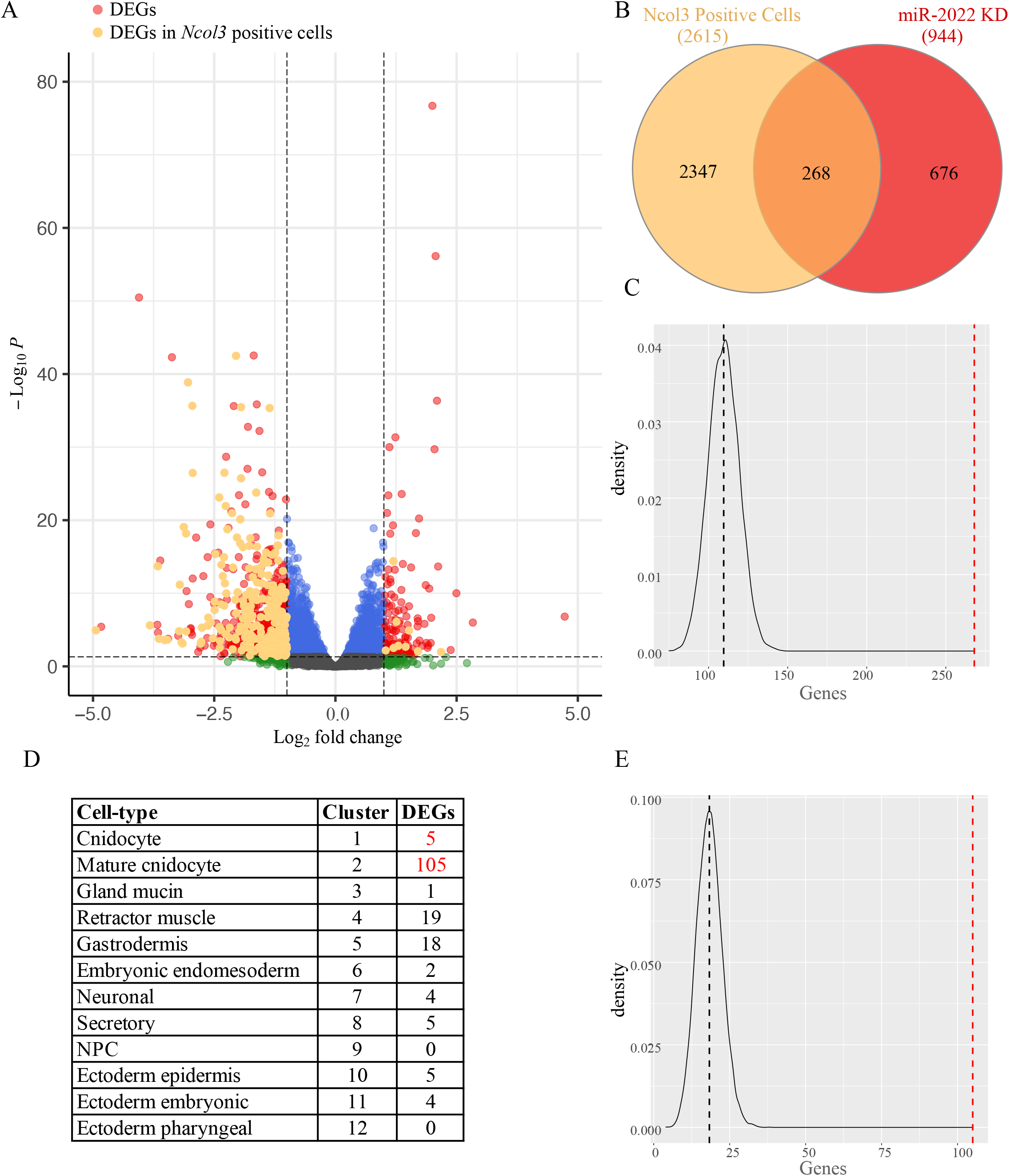
Transcriptome of *Nematostella* miR-2022 morphants. **(A)** Differentially expressed genes from 4 days old planulae injected either with control or Nv-miR-2022 MOs. Differentially expressed genes (red and orange) were defined by FDR < 0.05 and fold change ≥ 2. In orange: differentially expressed genes that are present in an independently obtained cnidocyte transcriptome (“Ncol3 positive cells”)^40^. (B) Overlapping portion of the differentially expressed genes identified in this study with genes from the cnidocyte transcriptomes^40^. (C) The likelihood of 268 cnidocyte specific genes overlapping with the list of 944 DEGs in miR-2022 morphants was assessed with Permutation test. 944 genes were randomly chosen 10,000 times (see methods for more details) and frequency of overlapping genes with genes of Ncol3 positive cells was plotted. **(D-E)** Differentially expressed genes in miR-2022 morphants that are found within cell type markers of the previously published single-cell expression data^19^, and permutation test. Black lines in **C** and **E** represent the expected overlap by chance relative to the observed (red line).

### Hyl1b is a plant-like miRNA biogenesis factor that neo-functionalized to control biogenesis of cnidocytes through controlling levels of Nv-miR-2022

Transcriptional regulators can be duplicated and then one of the paralogs can be recruited and neo-functionalized to facilitate the biogenesis of cnidocytes^28^. Previously we reported that the plant-like miRNA biogenesis factor Hyl1La, which is broadly expressed in *Nematostella*, regulates the biogenesis of multiple miRNAs in *Nematostella*, including miR-2022^11^. Here we focused on the paralog of Hyl1La, Hyl1Lb, which unlike the former is known to exhibit specific expression in cnidocytes^10^. We hypothesized that it could, therefore, be involved in the biogenesis of miR-2022. We designed a translation blocking MO to knockdown Hyl1Lb 4 days post injection of zygotes of wildtype females crossed to NvNcol3::memOrange2 males. Using stem-loop qPCR, we tested 5 different abundant miRNAs: we observed a significant strong reduction only for Nv-miR-2022 but not for Nv-miR-2025, Nv-miR-2026, Nv-miR-2027 and Nv-miR-2028 (**Figure 3A**). This suggests that Hyl1b is essential for the biogenesis of miR-2022 in cnidocytes, but not for the other tested miRNAs. Similarly to Nv-miR-2022 morphants, we observed a significant reduction of glowing cnidocytes in Hyl1b morphants (**Figure 3B**). This effect was quantified and validated by sorting and counting Ncol3::memOrange2 positive cells from Hyl1b MO and control injected animals (*t*-test. p = 0.0318, **Figure 3C**, and **Figure S3B**). Overall Hyl1Lb morphants mirrored the phenotype of Nv-miR-2022 morphants regarding its impact on cnidocytes, lowering their abundance without having a severe impact on the development (**Figure S3A**). This is in contrast to morphants of other miRNA biogenesis factors *in Nematostella* including Hyl1La that block morphogenesis^11,37,38^.

**Figure 3:**
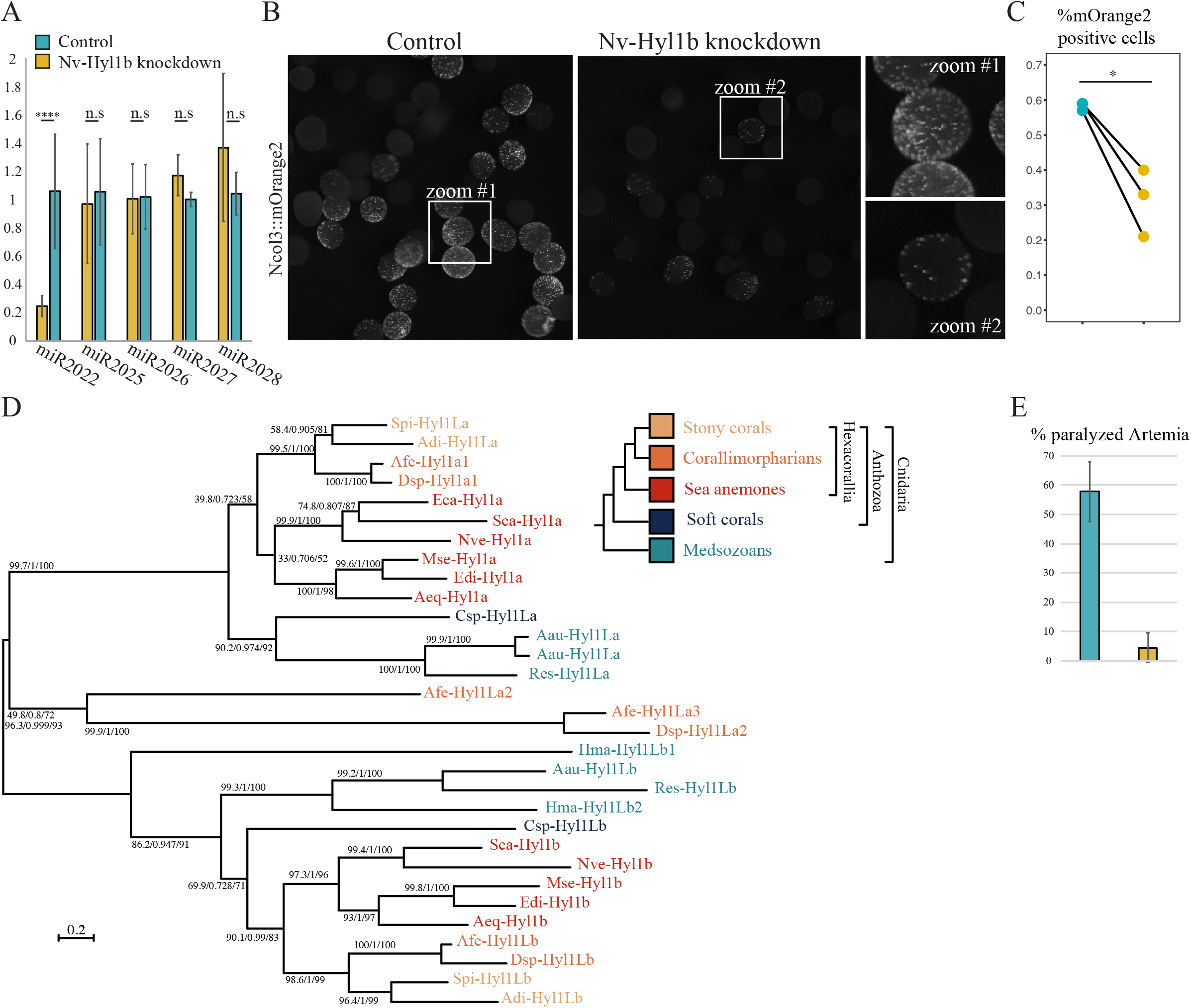
The impact of Nv-Hyl1b knockdown on cnidogenesis and prey paralysis potency in *Nematostella*. **(A)** Stem-loop qPCR assessment of Nv-miR-2022 levels as well levels of four additional non-cnidocyte miRNAs in *Nematostella* planulae 4 days post injection of control or Nv-Hyl1b MO into zygotes. Three independents biological replicates. (B) Intensity of mOrange2 in progeny of WT crossed with mOrange2::Ncol3 transgenes 4 days post injecting the zygotes with control or Nv-Hyl1b MO. Three independent biological replicates were subjected to flow cytometry analysis and the amounts of mOrange2 positive cells were quantified **(C). (D)** Phylogenetic relationship of cnidarian Hyl1 homologs. Topology represents maximum likelihood consensus phylogenetic tree. Support values of the SH-aLRT (50–100), approximate Bayes test (0.6–1.0), and ultrafast bootstrap replicates (50–100) appear from left to right near each relevant node. Aau, *Aurelia aurita*; Adi, *Acropora digitifera*; Aeq, *Actinia equina*; Afe, *Amplexidiscus fenestrafer*; Csp, *Clavularia*; Dsp, *Discosoma*; Eca, *Edwardsiella_carnea*; Hma, *Hydra magnipapillata* (vulgaris); Mse, *Metridium senile*; Nve, *Nematostella vectensis;* Res, *Rhopilema esculentum;* Sea, *Scolanthus callimorphus;* Spi, *Stylophora pistillata.* **(E)** Percentage of paralyzed Artemia 5.5 hours post incubation with control or Nv-Hyl1b MO 4 days old *Nematostella* planulae. Three independent biological replicates.

To better understand the relation between miR-2022 and Hyl1Lb, we collected the sequences of Hyl1 homologs from multiple cnidarian species and constructed a new phylogeny **(Figure 3D).** The grouping of some of the homologs from both Anthozoa (sea anemones and corals) and Scyphozoa (true jellyfish), two distantly-related cnidarian groups, with either Hyl1La or Hyl1Lb from *Nematostella* strongly suggests that Hyl1b originated through an ancient duplication and neo-functionalization event in the last common ancestor of all cnidarians.

Finally, as Hyl1Lb morphants mirror the reduction of cnidocytes through downregulation of Nv-miR-2022, we hypothesized that these morphants would exhibit defected capacity of predator paralysis similarly to Nv-miR-2022 morphants. Hence, we carried out the same paralysis assay presented in **Figure 1E** with Hyl1Lb 4 days old morphants. Similarly to the assays with Nv-miR-2022 morphants, *Artemia* nauplii incubated with control injected *Nematostella* planulae showed a significantly higher number of paralyzed/dead individuals compared to *Artemia* incubated with Hyl1Lb morphants (*t*-test, p=0.009112, **Figure 3E**, video **Data S3).**

Neo-functionalization of miRNA biogenesis/RISC proteins to specific tissues has been previously shown in bilaterians for Argonautes^41^, Dicers^42^, and other factors^43^. However, such events are mostly of recent origin and occur within distinct lineages of nematodes and vertebrates, not sharing an ancestral functional link to all bilaterians. In contrast, our phylogenetic analysis of Hyl1b strongly suggests that its recruitments to regulation of cnidogenesis occurred in the last cnidarian common ancestor **(Figure 3D).**

### miR-2022 is expressed in the stinging cells in cnidarian species separated by >600 million years

Using in-situ hybridization we tested if in addition to *NR2^9^*, other targets exhibit overlapping expression with miR-2022 in cnidocytes *of Nematostella* (**Figure 4A–B**). To this end, we generated probes against the primary sequence of Nv-miR-2022 (pri-Nv-miR-2022) as well as against two additional targets: *NVE16448* and *NVE16498* (their target sites are cleaved by Nv-miR-2022 according to degradome sequencing^9^). We found that both targets exhibited cnidocyte specific expression and overlapped with pri-Nv-miR-2022 across early *Nematostella* development (**Figure 4A–B**). While *NVE16498* lacks a known function, *NVE16448* encodes a putative arylsulfatase (also known as chondroitinsulfatase). It was previously shown that non-sulfated chondroitin structurally stabilizes cnidarian cnidocysts^44^, raising a possible link between this enzyme and cnidocyte biogenesis.

**Figure 4:**
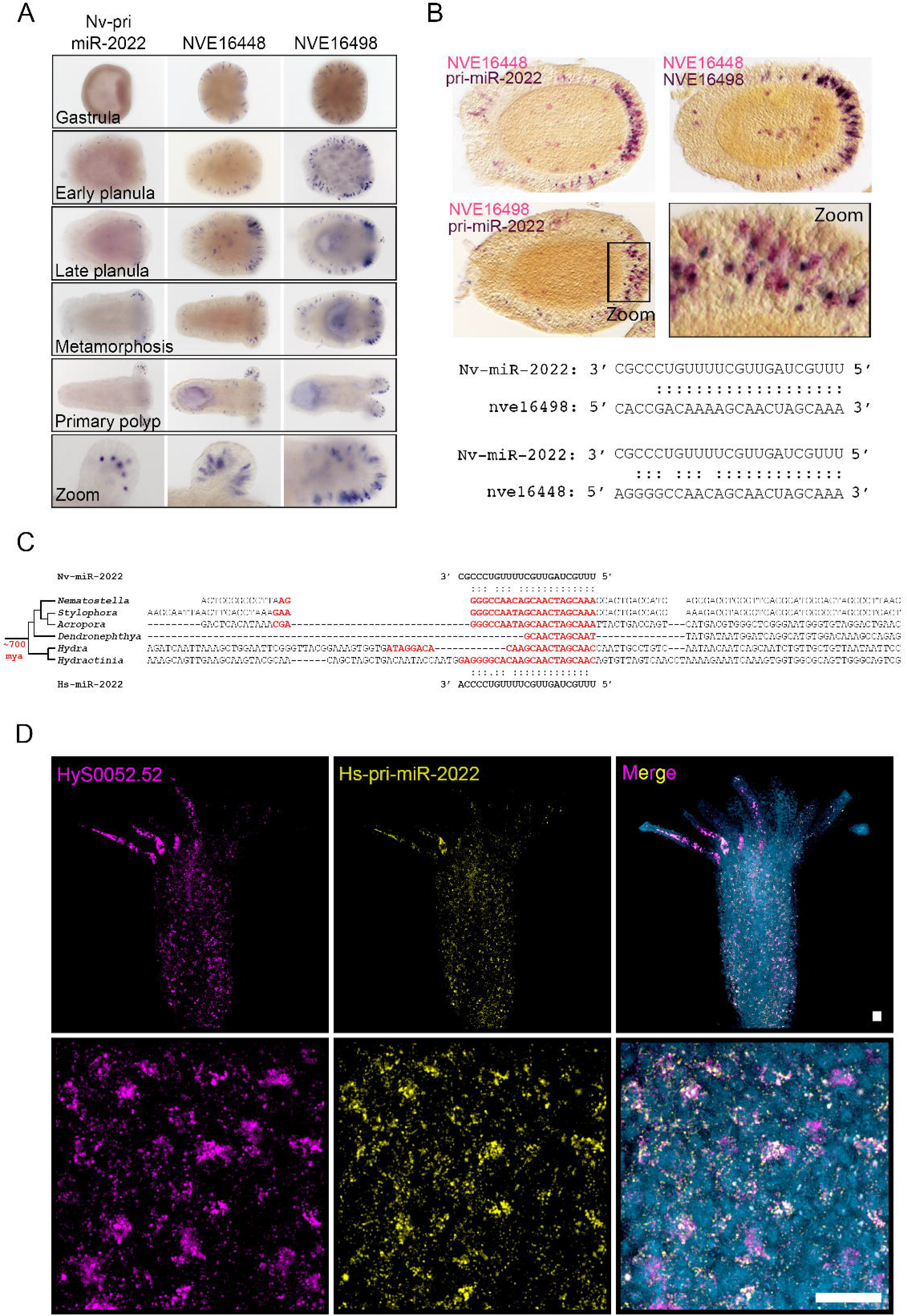
pri-miR-2022 with its targets in cnidocytes of *Nematostella* and *Hydractinia*. **(A)** Single ISH, NBT/BCIP staining using DIG-labelled probes for pri-miR-2022 and two of its targets across early developmental stages of *Nematostella.* **(B)** Double ISH, staining of pri-miR-2022 with its two targets in 4 days old *Nematostella* planulae. Samples were sliced into 14 μm sections to reduce background staining noise. In the two upper panels, DIG-labelled probes and NBT/BCIP staining were used for pri-miR-2022 staining, and FITC-labelled probes and FastRed staining were used for the targets. In the second from bottom panel, DIG-labelled probes and NBT/BCIP staining, and FITC-labelled probes with FastRed staining were used for visualization of nve16498 and nve16448 respectively. (C) Binding sites to miR-2022 on genes with top blast hits to nve16448 in distantly related cnidarians. **(D)** *Hydractinia* SABER FISH staining of Hs-pri-miR-2022 (yellow) and the predicted target Hys0052.52 (magenta).

Next, we computationally predicted targets of miR-2022 in several anthozoans (sea anemones and corals) and distantly related hydrozoan species. Surprisingly, a gene annotated as an arylsulfatase appeared in all species as one of the top targets. Reciprocal BLAST searches of this gene resulted in the predicted target of miR-2022 as the best hit in all species. This implies that homologs of NVE16448 are conserved targets of miR-2022 in Cnidaria (**Figure 4C**).

To further explore the functional basis for miR-2022 conservation, we used the publicly available single-cell RNA expression data *of Nematostella* and the hydrozoans *Hydra vulgaris* and *Hydractinia symbiolongicarpus*, which separated from Anthozoa at the dawn of cnidarian evolution. We found that the Arylsufatase-encoding gene with miR-2022 binding site (NVE16448, Hys0052.52 and t19037aep) was enriched in cnidocyte clusters of all three species (**Figure S4**). These results strongly suggest that regulation of NVE16448, Hys0052.52 and t19037aep represent an ancestral regulation of an Arylsulfatase-encoding gene by miR-2022 that persisted in extant cnidarians.

Finally, to test whether the expression of miR-2022 and its targets in cnidocytes is a cnidarian apomorphy, we characterized the expression patterns of Hys0052.52 and miR-2022 in *Hydractinia.* For this species a protocol for signal amplification by exchange reaction (SABER) fluorescence in-situ hybridization (FISH) has been optimized and enables the use of shorter probes^45^, which in our hands are essential for localizing the pri-miRNA in this species. Using double-FISH combinations for *Hydractinia* pri-mir-2022 (Hs-pri-miR-2022), Minicollagen 1 (Ncol1, a known cnidocyte marker^29,46^) and Hys0052.52, we found that pri-miR-2022 in this species co-localized with Hys0052.52 (**Figure 4D**) as well as with the cnidocyte marker, and that the target overlaps with Ncol1 (**Figure S4B**).

Precision of gene expression output is often difficult to control using transcription factors alone because transcription is an inherently noisy process^47^. Therefore, complex biological processes that govern development and cell differentiation in bilaterians recruited miRNA regulation to tune levels of dosage sensitive genes involved in these processes^48–50^. Two kinds of such miRNA-target modes of regulation are known: (1) coherent and (2) incoherent^51^. Coherent regulation can take part in biological processes such as promoting transitions of differentiating cells from their progenitors, by reciprocally inhibiting leftover transcripts belonging to the two opposing differentiated cell types^52^. In this type of regulation miRNAs show little to no overlap in expression with their targets^53^. In incoherent regulation, which is rare in bilaterians, miRNAs exhibit spatial-temporal overlap with their targets. It may seem puzzling to simultaneously activate gene transcription with its post-transcriptional repressor. However, this mechanism was shown to enable reaching precise expression levels by setting a threshold that prevents the target from exceeding a certain level, regardless of its transcriptional intensity, enabling to orchestrate precise stoichiometry of the targeted genes^54^. The co-expression of miR-2022 with several different targets in maturing cnidocytes of *Nematostella* and *Hydractinia* suggest that miR-2022 preserved its function of regulating cnidogenesis through incoherent regulation over 600 million years (**Figure 4, S4**).

To date, only a handful of pan-cnidarian transcriptional regulators essential for cnidocyte biogenesis are known. Among them, transcription factors such as SoxB2 act upstream to cnidogenesis and impact differentiation of both neurons and cnidocytes^17,45,55,56^, and PaxA which exhibits cnidocyte specific expression in Hydrozoa and Anthozoa^57,58^. The specific coexpression of miR-2022 with its targets in *Hydractinia* and *Nematostella* (**Figures 4**, and **S4**), as well as the single-cell RNA expression data supporting cnidocyte-specific genes carrying binding sites for miR-2022 in *Nematostella, Hydra* and *Hydractinia* strongly suggest that on top of transcriptional regulation, post-transcriptional regulation by the highly conserved miR-2022 emerged in the last common ancestor of Cnidaria and played a crucial role in enabling the innovation of these novel cell types through regulation of their biogenesis. To our knowledge, this is the first time the functional importance of an individual miRNA is demonstrated in a non-bilaterian animal.

## Summary

To summarize, the emergence of cnidocytes at the dawn of cnidarian evolution required the acquisition of novel genes, rewiring the function of existing ones, and the emergence of complex regulatory mechanisms to orchestrate the timing and levels of gene expression to allow the formation of the highly complex stinging organelles. As a result of their efficiency in prey/predator paralysis, and their early origin, it is plausible that survival of cnidarians became highly dependent on these cells, resulting in a selective pressure to preserve them in members of all cnidarian lineages^59^.

In this work we provide evidence that the evolution of the cnidarian-specific cell type, the cnidocyte, was accompanied by the emergence of miR-2022 as a novel regulator of its biogenesis. This was facilitated by rewiring of the miRNA biogenesis factor Hyl1, following gene duplication, to regulate the levels of miR-2022 in these cells. Therefore, cnidocyte biogenesis is one of the most ancient, conserved miRNA-regulated cellular processes in animals, known to date.

## Acknowledgments

The authors are grateful to Dr. Michal Bronstein and Ms. Adi Turjeman (The Center for Genomic Technologies of the Alexander Silberman Institute of Life Sciences, The Hebrew University) for their help with high-throughput sequencing. The authors would like to thank Dr. Uri Gat (The Hebrew University) for granting access and guidance using his lab equipment. Additionally, the authors would like to thank Dr. Grigory Genikhovich (University of Vienna) for material and protocol exchange during the early stages this project’s establishment. AF was supported by EMBO Short Term Fellowship 8417 and Long Term Fellowship 914-2021 during parts of this work. This work was funded by the European Research Council Starting Grant (CNIDARIAMICRORNA, 637456) and Consolidator Grant (AntiViralEvo, 863809) to Y.M.

## Methods

### In situ hybridizations (ISH and SABER-ISH)

For *Hydroctinia* polyps, Oligo probes were generated as described in Kishi et al., 2019 using hairpins 27 and 30^60^. Tissue fixation and dehydration followed by SABER-ISH were carried out as previously described^45,61^ using the following probes:

Hys0052.52 Hairpin 30:

miRTarg2.30.1 GGTGGTCGGTAAATACAATCTCTTCCTCCATCACCACTTTAATACTCTC

miRTarg2.30.3 TTGCTAGTTGCTTGTGCCCCTCCATTGGTATTGTCATTTAATACTCTC

miRTarg2.30.4 CTCCAGTGCTATCCGATGGAGAGTAAATAAGATCCTTGTCTTTTAATACTCTC

miRTarg 2.30.8 CCAATATCTCCAGCTTCGCCTCTCGTTTCTTCACCCTTTAATACTCTC

miRTarg2.30.10 CGATGTCCTGCAATCATTGCACTTGGAAGATCATTATGCTTTAATACTCTC

Hys-Mir2022 hairpin 27:

Pri-miR-2022.27.1 ACG CCG TC ATTTCTTCAC AAG TTTTCATTTG AACATCAG AATTTCATCATCAT

Hys-Ncol1 hairpin

30:

Ncol1.30.1 GACCAAGAGACATCCGAGGATAAATCGGGCTGCCATTTTAATACTCTC

Ncol1.30.2 TTCGTGTAAAGTTTTGGCGTTAGCCATGGCCACACCTTTAATACTCTC

Ncol1.30.3 TGGTGGTCCACATGGGTTAGCTTCACGTTTTAACATTTTAATACTCTC

Ncol1.30.4 CACATTGTTGCACACAGACTGGTGGGCATGGAGCTGTTTAATACTCTC

Hys-Ncol1 hairpin 27:

Ncol1.27.1 GACCAAGAGACATCCGAGGATAAATCGGGCTGCCATTTTCATCATCAT

Ncol1.27.2 TTCGTGTAAAGTTTTGGCGTTAGCCATGGCCACACCTTTCATCATCAT

Ncol1.27.3 TGGTGGTCCACATGGGTTAGCTTCACGTTTTAACATTTTCATCATCAT

Ncol1.27.4 CACATTGTTGCACACAGACTGGTGGGCATGGAGCTGTTTCATCATCAT

Colorimetric ISH and double ISH in *Nematostella* combining nitro-blue tetrazolium, 5-bromo-4-chloro-3’-indolyphosphate (NBT/BCIP) and FastRed (Roche, Germany) staining was performed according to established published protocols^62,63^ using the following DIG and FITC labeled probes:

Nv-pri-miR-2022:

GTAAATCAGCGTGGGGATGCAGACAGATAATCTCGGGATGAGTTCGCCTGAATGATCAAGATGAGTTCGCCTGAAAGTCGGGATGAGATCGCCTGAGAGTCGGGATAAGATCGCCTGAAAGTCGGGATAAATCAACTGTCAAGTGGTTGTCATTTGCTAGTTGCTTTTGTCCCGCCTTTTCTTTTTGTCCCGCCTTTTCTGCGAATTGATCACGTGATGTGACGTCATCACTGCCCAATTACAAATGCTGCAATATATTGAGGTACGTAAACGTTAGATGTTGCGCATATACCTCATACGCGTAATAACTCTGTGTAAATGTGAATGGTAGTAGAAAAAAATTAAATAAACCCAATT

Nve16498:

CCGTACCCATACCCATGCCTCATAATATGTCTACCACAACCAACACCACCTCCAGTAACCACACCCGAACCAGTAACCACACCCGAACCAGTAACCACACCCGAACCAGTAACCACGCAAAAGCCAGTAACGACGCCTAAGCCAACAACACCCAGTACAACCCCTAAACCTACAACTCCACCAACTCCTCCTTCACCTGTCAATGGTGGTTATACACCATGGACTGAATGGACCGAGTGCTCTGCCACGTGTGGGGGTGGTATCCAGCAAAGGACCAGAGCCTGTTCAAATCCTGCACCAAAGAACAATGGCACAAGCTGTGAATCACTAGGGCCATCATTTGAGACCCAAGCTTGCAATTCCAAACCATGTCCAGTTGATGGGGCTTACAGTGAGTGGAGTAAGTTTGACAAGTGTAGCAAATCTTGCGCTGGTGGAGTACAGATGCGGTCCAGAGAGTGTAACAACCCTGAGCCACAGTATGGCGGCAAGACTTGTAGCTATCTGGGGCCGGCCAACGAGACCAGAGCTTGCAACACTTTCTTCTGCCCAATTGATGGAGGTTACTGTGATTGGTCGGAATACATGTCATGTAGTGTGACGTGTGGGGGTGGTGTACAATACAAGACACGCACATGCACTAACCCTCCACCACAGCACGGAGGCAAGAACTGCTCTGAACTTGGGCCCTCGCGCATGTCACGTGACTGCAACACACACAGCTGTCCAGTTGATGGAGGCTACTCGCCTTACAGCAAATGGTCCGACTGTACAGAGACATGTGGTGGAGGTACACAGATTCGAACACGCACGTGCACTAACCCACCCCCTAAGCATGGTGGGCGCGACTGCTCACGGGACGGGCCAGACTTTGAGACTCGTGAATGCAACACTCAGGCTTGTCCAATCGACGGCGGTTTCACATCGTACACCAACTACTCCGAGTGCAGTAAAACATGTGGGGGCGGGACCCAGGTGCGCCTACGCTCGTGCACCAACCCTGAACCGCAGTACGGTGGCAAGAACTGCACGGGATCATACAAGCAAATTCG

Nve16448:

GATCTTGAATAACTACTACGTGTCTCCCATGGACACCCCGTCACGTGCGTCGTTCATGACGGGAAAGTACCCCATCCATATGGGTGTGCAACATGACACACTGCACAACAGGCAGCCCTTCGGGGTGCCGCTAACCGAGAAGTTTCTCCCGGAGTTTTTGAGAGAGATGGGCTACCAGACACATGCTGTCGGCAAGTGGCAGCTGGGGTTCTTCGCCAAGGAGTACACGCCCACCTATAGGGGATTTGACTCGTTCTTCGGCTTCTGGACCTCCCATGAAGATTACTACAATCACGTGGCCAATGACGGCGGCTACGGCATCGACCTCAGGAGGAACTTGGATGTCGTGTATAACGAGACAGGCGTATACGGTACGGAGCTTCTCGCTCGAGAAGCCGACGAAGTAATCGAGAATCATTCGGGAGACAAGCCACTGTTCCTGTACCTAGCACATCAGGCTGTGCACGTCGGGAATATGGACGAGCCGCTCCAGGCCCCTAAGCGTCACGTGGACAAGTTTAAGTACATCACGGATGAACGCAGAAGGACATTCGCCGGAATGGTATCCTCCCTTGACGAATCCATGCGCCAGCTTGTAACATCACTCAAACGCAAAGGCTTGTACCAAAACTCCATAATTATCTTCACCACTGACAACGGTGGCGCCGCTGGTGGACTAGACATGAGCGCGGGCTCAAACTTTCCCCTGCGCGGTAACAAGAACACCCTGTGGGAGGGTGGGGTGCGTGGTGTGGCATTTGTCCACAGTCCTCTTATCAAG

### *Nematostella* culture

*Nematostella* polyps were grown in 16%o sea salt water at 18 °C. Polyps were fed with *Artemia salina* nauplii three times a week. Spawning of gametes and fertilization were performed according to a published protocol^64^. In brief, temperature was raised to 25 °C for 9 h and the animals were exposed to strong white light. Three hours after the induction, oocytes were mixed with sperm to allow fertilization.

### Nv-miR-2022 and Nv-Hyl1b knockdowns

Male adults of the Ncol3::memOrange2 transgenic line (generated in a previous study^28^) were spawned as previously described^64^ and crossed with WT eggs. The gelatinous sack surrounding the eggs was removed using 4% L-Cysteine (Merck Milipore, USA) and followed by microinjecting the zygotes with antisense MOs. Next, zygotes were cultured at 22 °C in 16%o artificial seawater in the dark. The MO sequences were designed and synthesized by Gene Tools, LLC (USA). miR-2022 MO sequence: ACCACTTGACAGTTGATTTATCCCG.

Standard control MO sequence: CCTCTTACCTCAGTTACAATTTATA.

Hylb translation blocking MO: CTGATCCTGGATGTTGAAACCTCAT.

miR-2022 MOs (0.9mM), Hyl1b MO (0.45mM) and standard control MOs at equal concentrations were injected the same day into zygotes from the same batch to generate one biological replicate. In total, three biological replicates of ~600 injected animals each were generated for miR-2022 and control MO and subjected to morphological characterization, flow cytometry analysis and paralysis assays.

### RNA sequencing

Total RNA was extracted using TRIzol Reagent (Thermo Fisher Scientific) and following manufacturer’s protocol. Samples were treated with 2 μl of Turbo DNAse (Thermo Fisher Scientific) and undergoing an additional round of extraction using TRIzol Reagent. The total RNA quality was assessed using Bioanalyzer Nanochip (Agilent) with all samples having RNA Integrity Number (RIN) > 7.

### Stem-loop qPCR

For the quantification of miRNAs, we used stem-loop primers^35^ for miR-2022, miR-2025, miR-2026, miR-2027 and miR-2028 from our previous study^11^. For cDNA preparation, 100 ng of total RNA was reverse transcribed using the Superscript III Reverse Transcriptase (Thermo Fisher Scientific). Specificity of miRNA primers was determined with end point PCR^65^. For this, we used 2 μl of cDNA as template, miRNAs-specific forward primer and stem-loop-specific reverse primer and run the PCR at 94°C for 2 min, followed by 35 cycles of 94°C for 15 s and 60°C for 1 min. For differential expression analysis, we ran qRT-PCR with 5sRNA as an internal control. For all the real-time experiments, we used Fast SYBR Green Master Mix (Thermo Fisher Scientific) and samples were run on StepOnePlus Real-Time PCR System (Thermo Fisher Scientific). Experiments were performed in three independent biological replicates and two technical replicates and data was analyzed using 2^-ΔΔCt^ method^66^.

### Paralysis assay

4 days post injection with control, miR-2022 or HYIlb MOs, *Nematostella* planulae were incubated with *Artemia* in 0.5 ml *Nematostella* medium in 24 well plates. In each biological replicate ~60 planulae were divided into 3 wells (20±3 per well) and incubated with 8-10 *Artemia* per well. Swimming *Artemia* were documented at time 0 and 5.5 hours from the start of incubation. As a negative control, *Artemia* were incubated in *Nematostella* medium without planulae.

### Planulae dissociation and flow cytometry analysis

Animals were dissociated into single cells as previously described^67^. Briefly, 4 days old planulae were placed in 1/3 strength calcium/magnesium free artificial seawater (17 mM of TrisHCl, 165 mM of NaCl, 3.3 mL of KCl and 9 mM of NaHCO3; final solution pH 8.0). Next, planulae were washed twice with 1/3 strength artificial seawater and incubated with 50 μg/mL liberaseTM (Roche) at 37 °C for 10–20 min with occasional pipetting, until fully dissociated. About 200-500 individuals were used per tube. The reaction was stopped by adding 1/10 volume of 500 mM EDTA solution. Cells were centrifuged at 500× *g* at 4 °C and resuspended in 1/3 strength artificial seawater containing 2 μg/mL calcein AM (Enzo) and 100 nM sytox blue (BioLegend) to monitor viability. The suspension was filtered using 5 mL round-bottom tubes with 35 μm cell strainer cap (Corning). Cells were incubated with the viability dyes for 20 min at room temperature and examined by fluorescence microscopy or flow cytometry.

For miR-2022 knockdowns, FACSAria III (BD Biosciences) equipped with 488-, 405- and 561-nm lasers was used to determine memOrange2 expression in single cells. Per run, 100,000 events were recorded. Sytox blue positive events were excluded and calceinAM positive events were included. FCS files were further analyzed using FlowJo V10 (BD Biosciences). For Hylb knockdowns, Cellstream (Merck) sorter was used using the same lasers, and 50,000 event were recorded.

### Prediction of miR-2022 targets

Putative miRNA targets in *Hydra* and *Hydractinia* were identified with psRNATarget^68^ with the following parameters: Maximum expectation score 2, GU penalty 0.5, extra weight seed 1.5, mismatches in seed 2, other mismatches 1, penalty for opening a gap 2, penalty for extending the gap 0.5, HSP size 19, seed region 2-13.

### RNA sequencing analysis

Raw reads were trimmed and quality filtered by Trimmomatic^69^, using previously published parameters^70^. Reads were mapped to a modified *Nematostella*^71^ which included an additional contig that encodes for memOrange2. Mapping was performed using STAR^72^ and the gene counts quantified using RSEM^73^. Differential expression analyses were performed using scripts from Trinity^74^ two both DESeq v2.1^39^ and edgeR v3.16^75^. Gene models used in all downstream analyses were from previously published annotations (Schwaiger et al., 2014^76^; https://figshare.com/articles/Nematostella_vectensis_transcriptome_and_gene_models_v2_0/807696). Differentially expressed genes were defined by FDR < 0.05 and fold change ≥ 2. Genes identified both methods were considered as differentially expressed. Biological replicates were quality checked for batch effect using sample correlation and principal component analysis. Volcano plots were generated using EnhancedVolcano (https://github.com/kevinblighe/EnhancedVolcano).

We then investigated the impact of miR-2022 knockdown on the transcriptional profile of nematocytes. To do this we performed comparative analysis using previously published RNAseq data^40^, generated from the primary polyps of a nematocyte transgenic reporter line, NvNcol3::mOrange2^28^. Raw reads were downloaded from the Sequence read archive (BioProject: PRJEB40304) for both NvNcol3::mOrange2 positive and negative cells and DGE analysis performed as previously described. To gain further insights into the expression of the targets of miR-2022 and other genes of interest at the cellular resolution, we investigated new scRNA-seq data across a time course of *Nematostella* development that was recently published Steger, J. *et al 2022^19^*. Counts matrix and cell barcodes were downloaded from the Gene expression omnibus (accession: GSE200198) and scRNA-seq was performed using previously published methods (https://github.com/technau/CellReports2022/blob/main/DataS6_Merge2Figures.R) We further tested whether genes significantly affected by miR-2022 KD are enriched in NvNcol3::mOrange2 positive cells as well as any specific cell cluster from scRNA-seq data across the development of *Nematostella.* To do this we performed permutation tests to validate that the overlap between genes differentially expressed following miR-2022 KD and those upregulated in NvNcol3::mOrange2 positive cells or marker genes of cell clusters are significant. Genes that had < one count per million for at least two libraries from the miR-2022 KD RNAseq data were removed to generate a background dataset. From this background, 944 genes were randomly selected 10,000 times and the overlap to genes upregulated in NvNcol3::mOrange2 positive cells and marker genes of cell clusters was compared.

### Phylogenetic analysis

To construct the phylogenetic tree we selected representatives of HYL-1-Like sequences from anthozoans and medusozoans available in public databases. Specifically, *Nematostella* Hyl1a/b amino acid and nucleotide sequences were BLAST searched against protein and nucleotide databases of species present in the Reefgenomics database (*Acropora digitifera, Actinia equina, Amplexidiscus fenestrafer, Discosoma, Hydra magnipapillata (vulgaris), Stylophora pistillata)^77^*. The sequences Hyl1 from the rest of the species in the phylogeny (*Aurelia aurita, Clavularia, Edwardsiella_carnea, Metridium senile, Rhopilema esculentum, Scolanthus callimorphus*) are top BLAST hits against either Transcriptome Shotgun Assembly or RefSeq^78^ databases of NCBI. The Hyl1s amino acid sequences were aligned using MUSCLE^79^and low quality alignment regions were removed by TrimAl^80^ using the - automatic1 for heuristic model selection. The Maximum-Likelihood phylogenetic trees were constructed using IQ-Tree^81^ with the VT+R3 model which was the best fitting model according to the Bayesian information criterion (BIC). Support values of the ML tree were calculated by three different methods: 1,000 ultrafast bootstrap replicates^82^, 1,000 replicates of the Shimodaira-Hasegawa approximate likelihood ratio test (SH-aLRT) and an approximate Bayes test.

## Figures

**Figure S1: The impact of miR-2022 knockdown on cnidogenesis and development of *Nematostella*. (A)** Nv-miR-2022 precursor with highlighted region targeted by miR-2022 MO. (B) Dissociated cells of control and miR-2022 MO injected planulae 4 days post injection. Arrowheads mark cnidocytes. (C) Flow cytometry analysis of mOrange2::Ncol3 positive cells control and miR-2022 MO injected transgenes. Y axis represents the intensity of mOrange2 and cell viability is measured using CaleinAM on the X axis. Wildtype animals (right panel) were used set the threshold of the mOrange2 positive gate. Percentages of positive cells are indicates within the gates on the plot. (D) Cells from positive and negative gates were collected and visualized under a microscope to further confirm that glowing cells from the positive gates correspond to cnidocytes. (E) Intensity of mOrange2 and morphology of WT crossed with mOrange2::Ncol3 progeny 9 days post injecting the zygotes with control or miR-2022 MO. (F) Analysis of *Nematostella* single-cell RNA sequencing data generated by Steger, J. *et al.^19^* Arrowhead indicates cluster of cnidocytes and color intensity corresponds to expression intensities of NvAGO1.

**Figure S2:** differential expression analysis to Identify cnidocyte specific genes in *Nematostella* primary polyps of Ncol3::mOrange2 transgenic line (see methods for more details).

**Figure S3: The impact of Nv-Hyl1b knockdown on cnidogenesis and development of *Nematostella*. (A)** Intensity of mOrange2 and morphology of WT crossed with mOrange2::Ncol3 progeny 9 days post injecting the zygotes with control or Nv-Hyl1b MO. **(B)** Flow cytometry analysis of mOrange2::Ncol3 positive cells control and Nv-Hyl1b MO injected transgenes. Y axis represents the intensity of mOrange2 and cell viability is measured using CaleinAM on the X axis. Wildtype animals (right panel) were used set the threshold of the memOrange2 positive gate. Percentages of positive cells are indicates within the gates on the plot.

**Figure S4: Expression of miR-2022 predicted targets in cnidocytes of *Hydra* and *Hydractinia*. (A)** Single cells expressing NVE16448, Hys0052.52 and t19037aep in *Nematostella, Hydractinia* and *Hydra* (respectively). In *Hydractinia* expression intensities of Ncol1 and Hys0052.52 enable the visualization of the cnidocyte cluster of cells (bottom left) and the intensity of overlapping expression of Hys0052.52 within this cluster. In *Nematostella* the plot was generated as described in methods. In *Hydractinia* and *Hydra* plots were generated using the Single Cell Browser within the *Hydractinia* Genome Project Portal (https://research.nhgri.nih.gov/hydractinia) and using the *Hydra* single cell portal^58^ **(B)** SABER FISH staining of Ncol1 (green) with Hys0052.52 (magenta) or with Hs-miR-2022 (yellow) in *Hydractinia* polyps.

