## Supplementary figures and images for "A pan-cnidarian microRNA is an ancient biogenesis regulator of stinging cells"

### Supplementary Figure S1

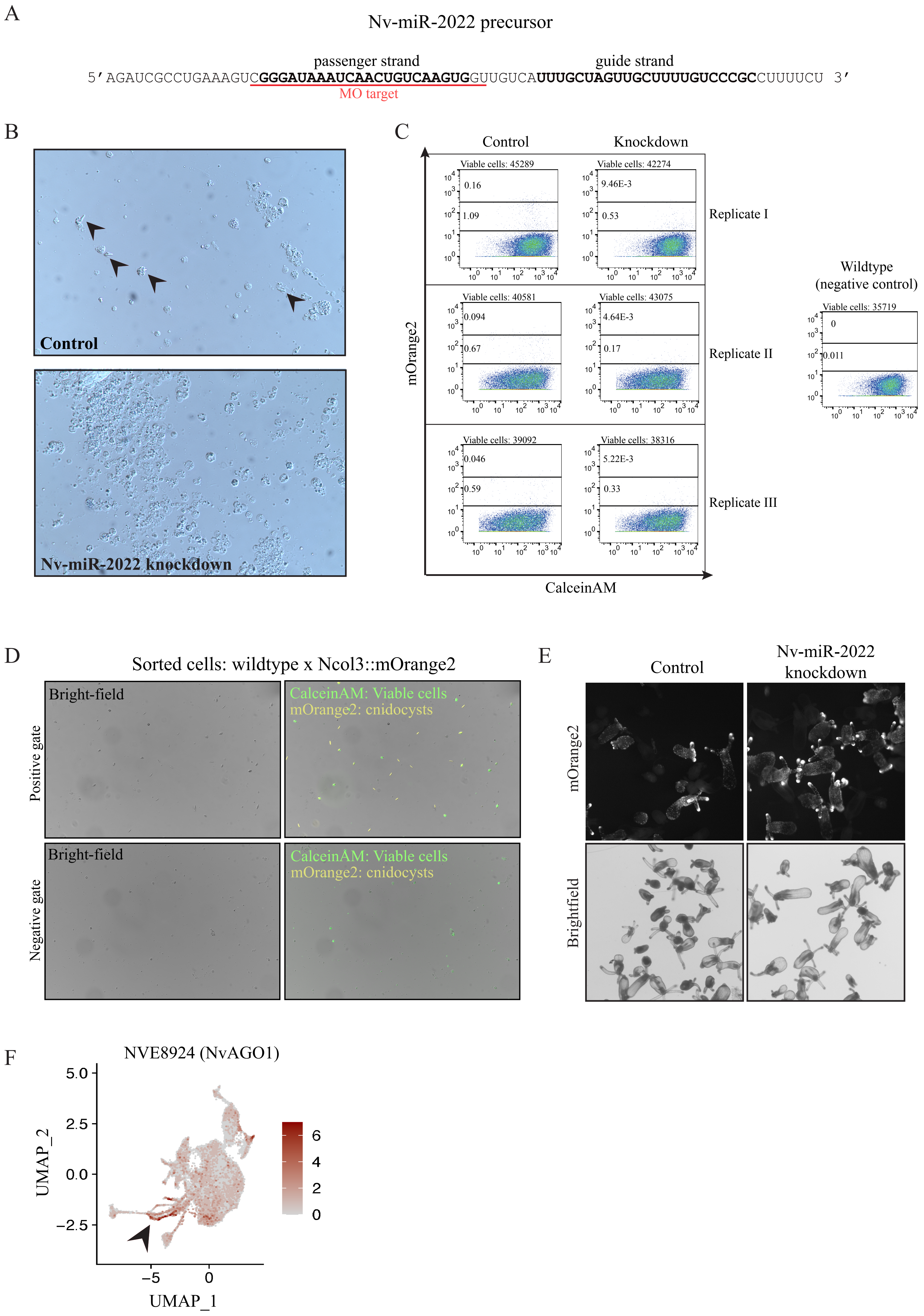

### Supplementary Figure S2

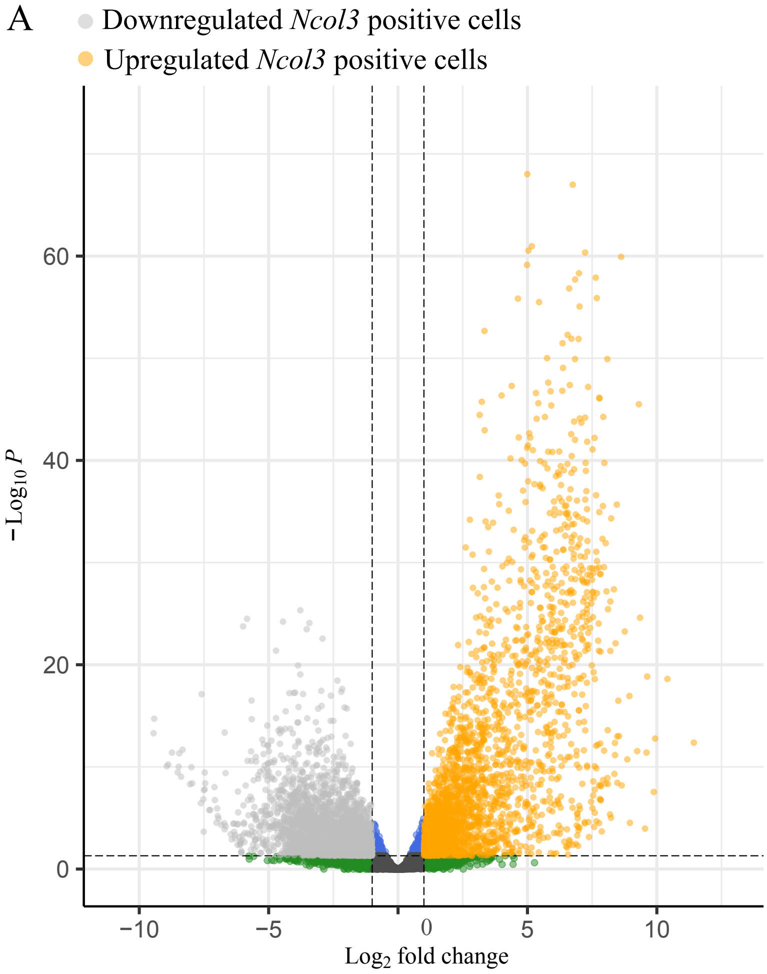

### Supplementary Figure S3

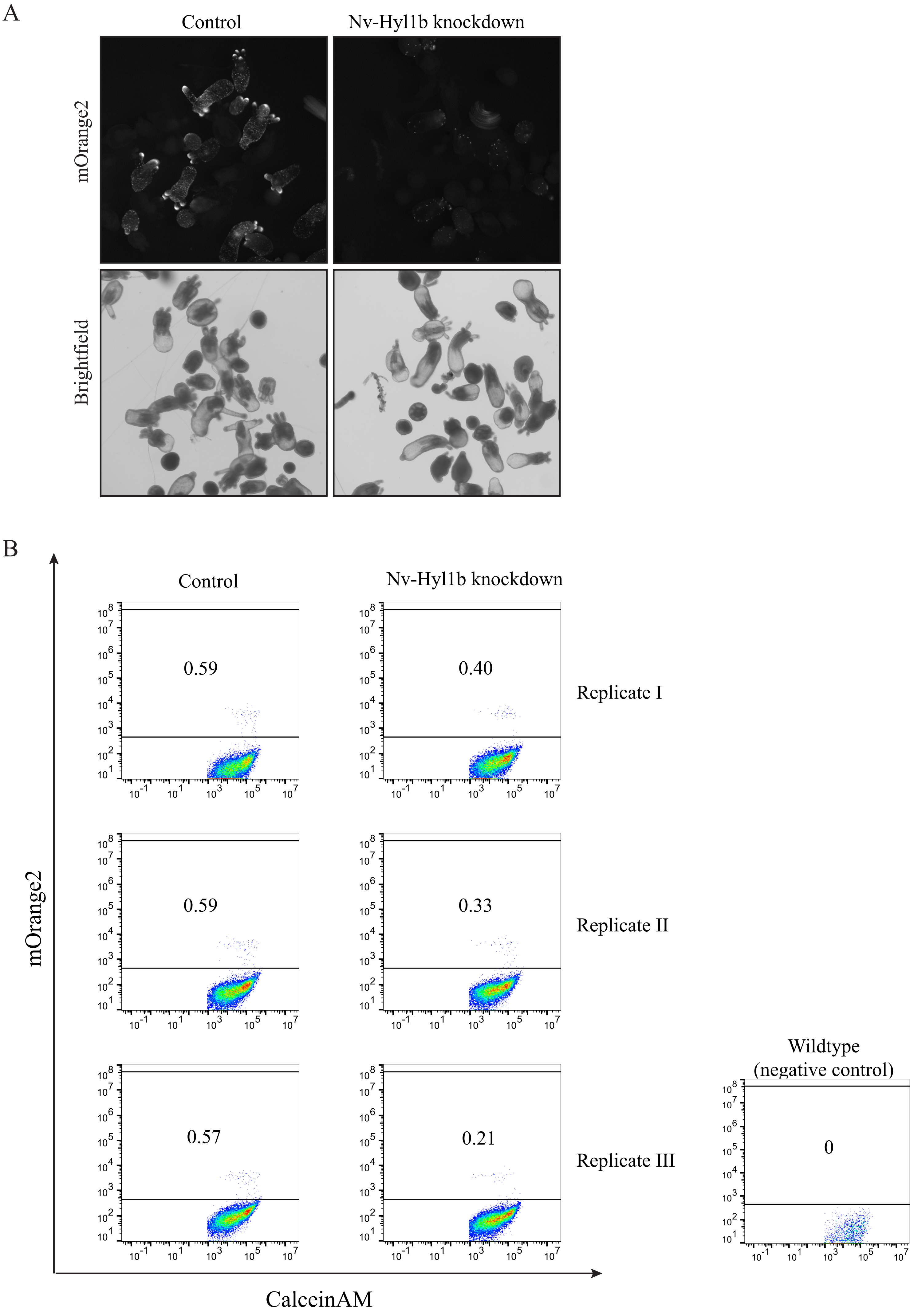

### Supplementary Figure S4

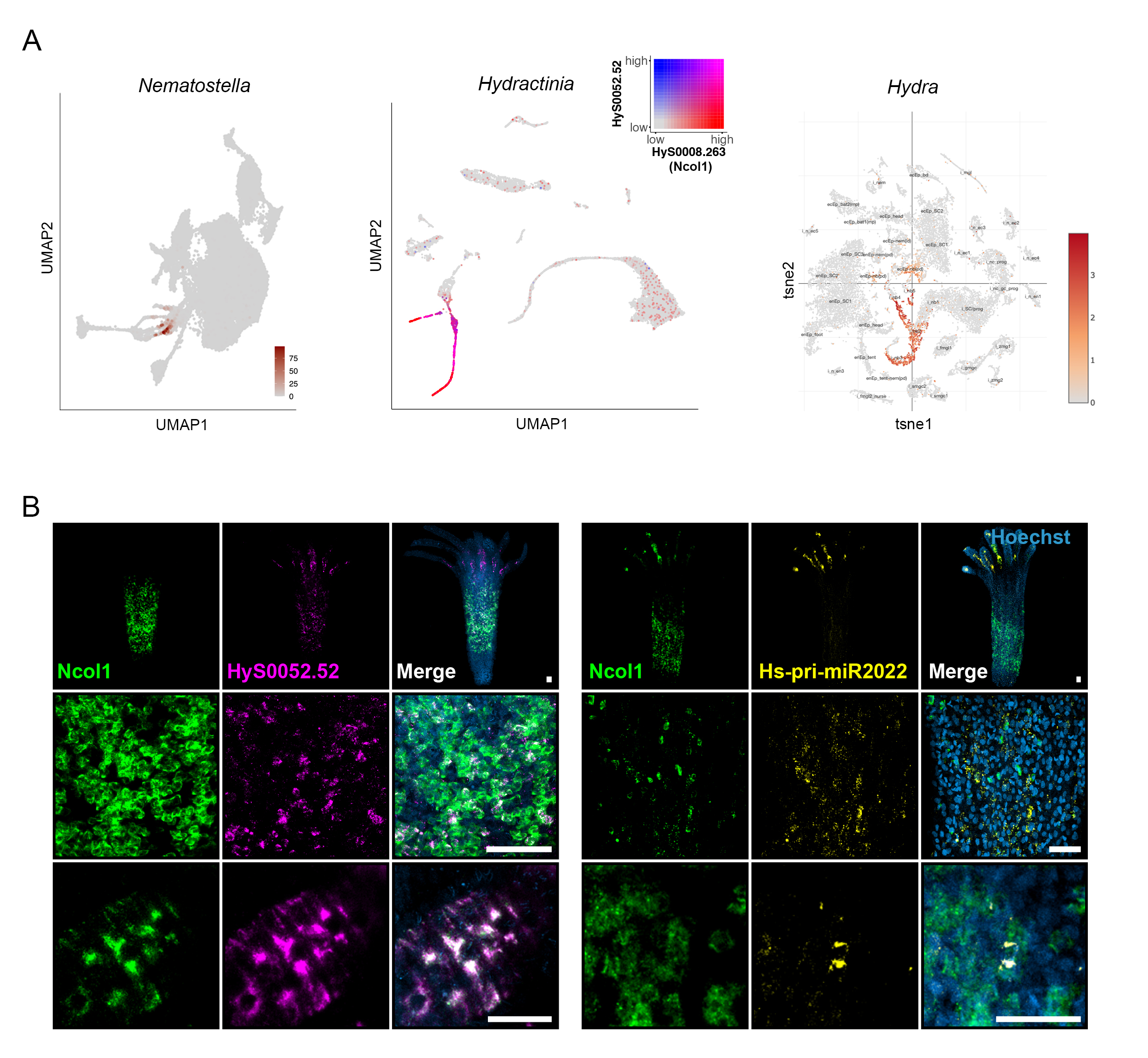
